# A Human iPSC-derived Organoid System to Model Myelin Destruction and Repair

**DOI:** 10.1101/2024.05.19.594912

**Authors:** Shwathy Ramesan, Joel Mason, Molly E.V. Swanson, Samuel Mills, Bradley J. Turner, Matthew Rutar, David G. Gonsalvez, Samantha K. Barton

## Abstract

We engineered a novel human iPSC-derived organoid system enriched with mature, myelinating oligodendrocytes and functionally reactive microglia. This model enables the study of human CNS remyelination following a demyelinating insult, encapsulating: myelin fragmentation, microglial clearance of myelin debris and oligodendrocyte genesis, differentiation, and axon ensheathment. This system provides a powerful and unparalleled capacity to interrogate complex human cell behaviour, relevant to those studying glial biology and neurological disease mechanisms.

## Main Text

Axons in the CNS are ensheathed by myelin, a specialised lipid membrane, that is generated by oligodendrocytes. Both myelin and oligodendrocytes can be destroyed in CNS injury, which can lead to debilitating functional deficits and irreversible neurodegeneration. Fortunately, the CNS has some capacity to generate new myelin membranes as part of a repair process called re-myelination. At present, we can only study these dynamic processes of remyelination in animal models. Although conserved molecular mechanisms have been identified between human myelin biology and that of non-human animal models, distinctions in cell behaviours are evident [1]. Using iPSC-derived organoids, we aimed to generate an organoid system that modelled human myelin damage and repair to generate a platform that could potentially be used for identifying therapies that successfully promote oligodendrocyte regeneration and myelin repair.

Our first step was to successfully engineer iPSC-derived CNS organoids containing myelinating oligodendrocytes and functional microglia (Suppl Fig 1a, b). The host organoids were spinal cord patterned organoids enriched with extensive and mature myelinating oligodendrocytes [2, 3]. *In vivo*, microglial/macrophage precursors migrate into the developing CNS after neurogenesis commences [4] so we seeded separately differentiated iPSC-derived macrophage-like precursor cells into organoids to more closely model CNS development. The macrophage-like cells were differentiated separately from iPSCs [5] and seeded into organoids three weeks into the maturation phase of culturing. Through intrinsic organoid signalling, the macrophage-like cells successfully differentiated to microglia by 11 weeks of maturation, evidenced by a reduction in CD45+ expression and enriched P2YR12+ and CX3CR1+ expression (Suppl Fig 1 a-e). Live imaging of their tdTomato fluorescent reporter showed successful integration into the organoids, mimicking the colonisation seen in human development (Suppl Movie 1), and they were dynamically reactive to stimuli implying functionality (Suppl Movie 2). Our organoid system thereby presented an ideal base for aiming to interrogate human myelin biology.

Myelin perturbations underpin a plethora of neurological diseases; however, a prevalent disease in need of strategies to promote myelin repair is Multiple Sclerosis (MS). MS is characterised by autoimmune attacks targeting myelin and the oligodendrocytes that produce it. Myelin and oligodendrocyte loss leads to irreversible neural degeneration and disease progression. Currently, no treatment options are available to MS patients that can promote remyelination, despite success in pre-clinical animal models, highlighting the essential need for a human model system that lends itself to identifying mechanisms, and screening drugs, to promote myelin repair. In order to trigger demyelination in our newly developed iPSC-derived organoids containing myelin and microglia, we utilised lysolecithin (LPC), which disrupts lipid structure and causes focal demyelination in animal models of MS [6]. Organoids were subjected to 0.1 mg/mL LPC (or vehicle v/v; Suppl Fig 2a) to trigger demyelination (but not overt microglial activation; Suppl Movie 3, 4) for 16 h after which LPC was removed equating to time point 0. Organoids reproducibly showed significantly shorter myelin sheath length at all time points (Fig 1a, b) implying successful myelin fragmentation and therefore demyelination of our organoids. This phenotype was replicated using a second iPSC line (Suppl Fig 2c, d).

**Figure 1:**
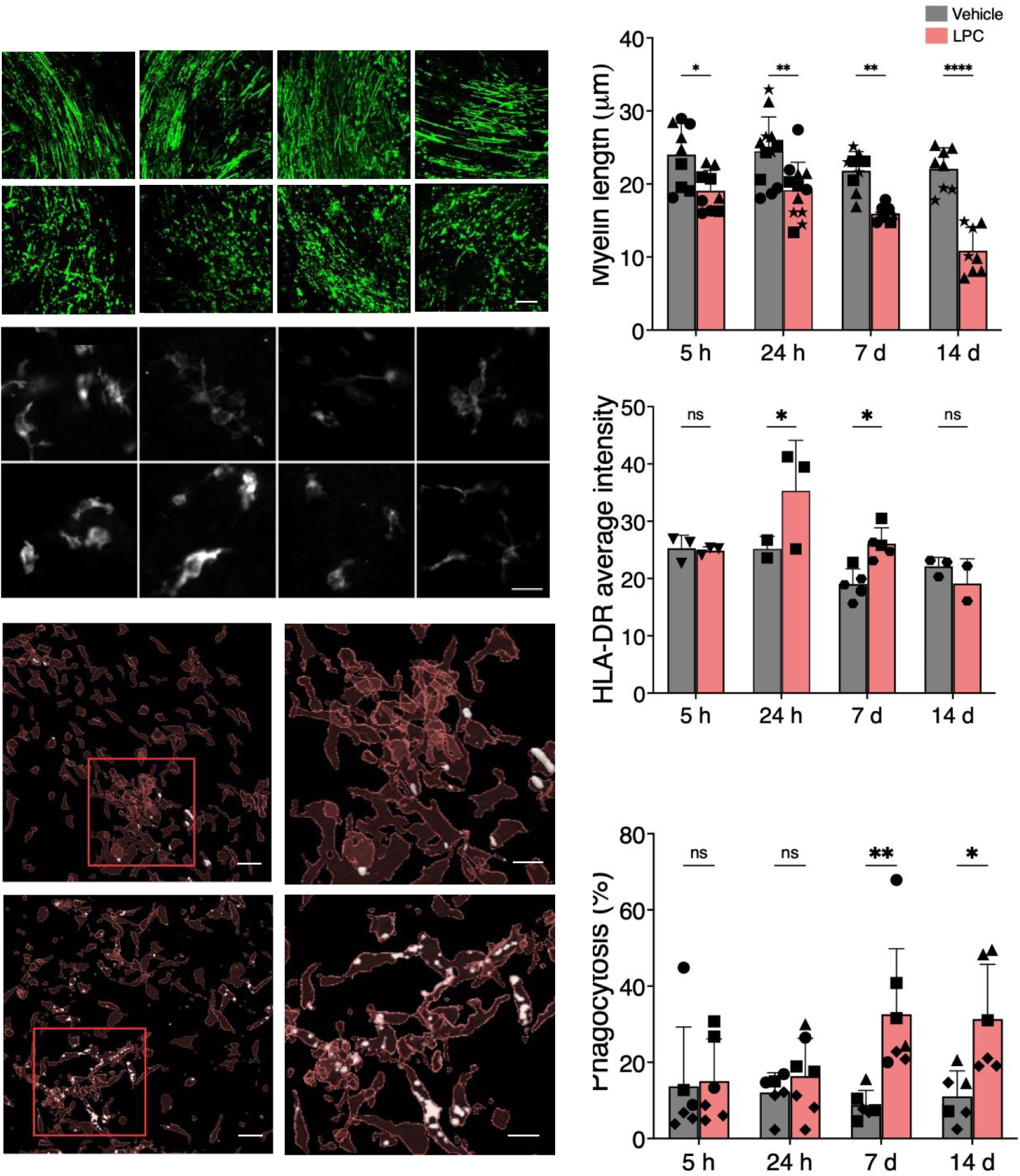
(a) LPC triggered overt MBP-positive myelin fragmentation comparable to Vehicle treated organoids at all time points; scale bar = 50 μm. (b) Average myelin length per organoid was significantly shorter in LPC treated organoids compared to Vehicle treated at all time points. (c) Iba-1-positive microglia expressed HLA-DR at all time points; scale bar = 15 μm and (d) expression was significantly higher 24 h and 7 d after LPC removal compared to Vehicle. (e) There was more extensive MBP-positive myelin debris within Iba-1-positive microglia; scale bar = 50 μm (20 μm in inset image highlighted by red box) and (f) significant phagocytosis was noted at 7 d and 14 d after LPC removal. Each data point depicts the average from an individual organoid; different symbols depict independent differentiations from iPSC. Data graphed as mean ± SD and compared using a 2-way ANOVA. ns = not significant, * p<0.05 ** p<0.01 *** p<0.0005 **** p<0.0001

Microglia play important roles in demyelination, myelin clearance and myelin regeneration [reviewed in 7]. Live imaging of microglia 7 d after LPC/Vehicle highlights the variability in morphology amongst the population (Suppl Video 5, 6) and quantification of the microglial morphology showed no consistent change to mean soma area (Suppl Fig 3a) or mean filament area (Suppl Fig 3b), as a surrogate for cell body and process area, respectively, between groups at any time point. To account for the heterogeneity, microglia were further stratified using multiplexed fluorescent immunolabelling (Suppl Fig 3c). Across Iba-1 positive microglia, the average tissue-wide expression of HLA-DR was significantly increased 24 h and 7 d after LPC removal compared to Vehicle (Fig 1c-d). HLA-DR is a marker of microglia “activation” and is upregulated during antigen presentation; its expression is increased in MS cases [8]. Distribution plots of microglia from Vehicle- and LPC-treated organoids were unimodal suggesting all microglia were increasing HLA-DR expression (rather than the emergence of an HLA-DR^high^ microglia population; Suppl Fig 3d). Expression of Iba-1, CD68, and L-ferritin also increased at 24 h and 7 d with reduction by 14 d (Suppl Fig 4). This led us to question whether microglia in our organoids have the functional capacity to clear myelin debris. There was a significant increase in co-localisation of MBP signal within Iba-1 positive microglia (Fig 1e) in LPC treated organoids after 7 d and 14 d, comparative to Vehicle (Fig 1f), suggestive of myelin debris phagocytosis by microglia. The co-localisation data were reproduced in organoids seeded with macrophage-like cells generated from the matched iPSC control line to the organoids as validation (Suppl Fig 5a-b).

Irrefutable evidence from animal studies shows that remyelination can take place by either: 1) existing oligodendrocytes that survive the demyelinating injury, or 2) by newly generated oligodendrocytes that produce new myelin [9, 10, 11]. In humans, however, from evidence using ^14^C isotope birthdating, it remains unclear whether oligodendrocyte precursors have the capacity to proliferate, differentiate and establish new myelin subsequent to a demyelinating injury in MS [1]. The more fundamental issue is knowing with absolute certainty that a remyelinating event is even occurring in human MS [12]. Unlike in animal models, in human tissue we lack the capacity to elicit demyelinaton and observe the ensuing remyelination by human oligodendroglia. Our model overcomes this hurdle, and enables us to observe the potential of human oligodendroglia to respond to demyelination. To distinguish between myelinating oligodendrocytes that survived the LPC insult and new cells generated from precursor cell proliferation and differentiation, the culture media was supplemented with the thymidine analogue ethynyl deoxyuridine (EdU) from the time of LPC removal (Fig 2a, Suppl Fig 2b). In LPC treated organoids, there was a significant increase in newly generated oligodendrocytes as a proportion of all oligodendrocytes (Fig 2b; p<0.05 for all). When assaying the proportion of newly generated oligodendrocytes that had formed myelin, there was a significant increase 8 w (p=0.0002) and 12 w (p=0.035) after LPC removal (Fig 2c). These data therefore allow us to conclude that human OPCs have the capacity to proliferate, differentiate and myelinate in response to a demyelinating insult. It is currently not possible in our system to assay oligodendrocytes that survive demyelination or to measure remyelination, per se. The applicability of this model system would be to identify therapeutic candidates that can successfully promote myelin repair and regeneration. Upon repeat of the remyelination assay alongside culturing organoids treated with clemastine fumarate (1 μM), the first efficacious remyelinating drug [13] to enter clinical trial for MS [14], a significant increase in new myelin formation was found comparative to LPC treated organoids cultured in a media devoid of growth factors, with no changes observed in any ethanol treated organoids (Fig 2d), highlighting the exciting potential of using this model system for drug screening.

**Figure 2:**
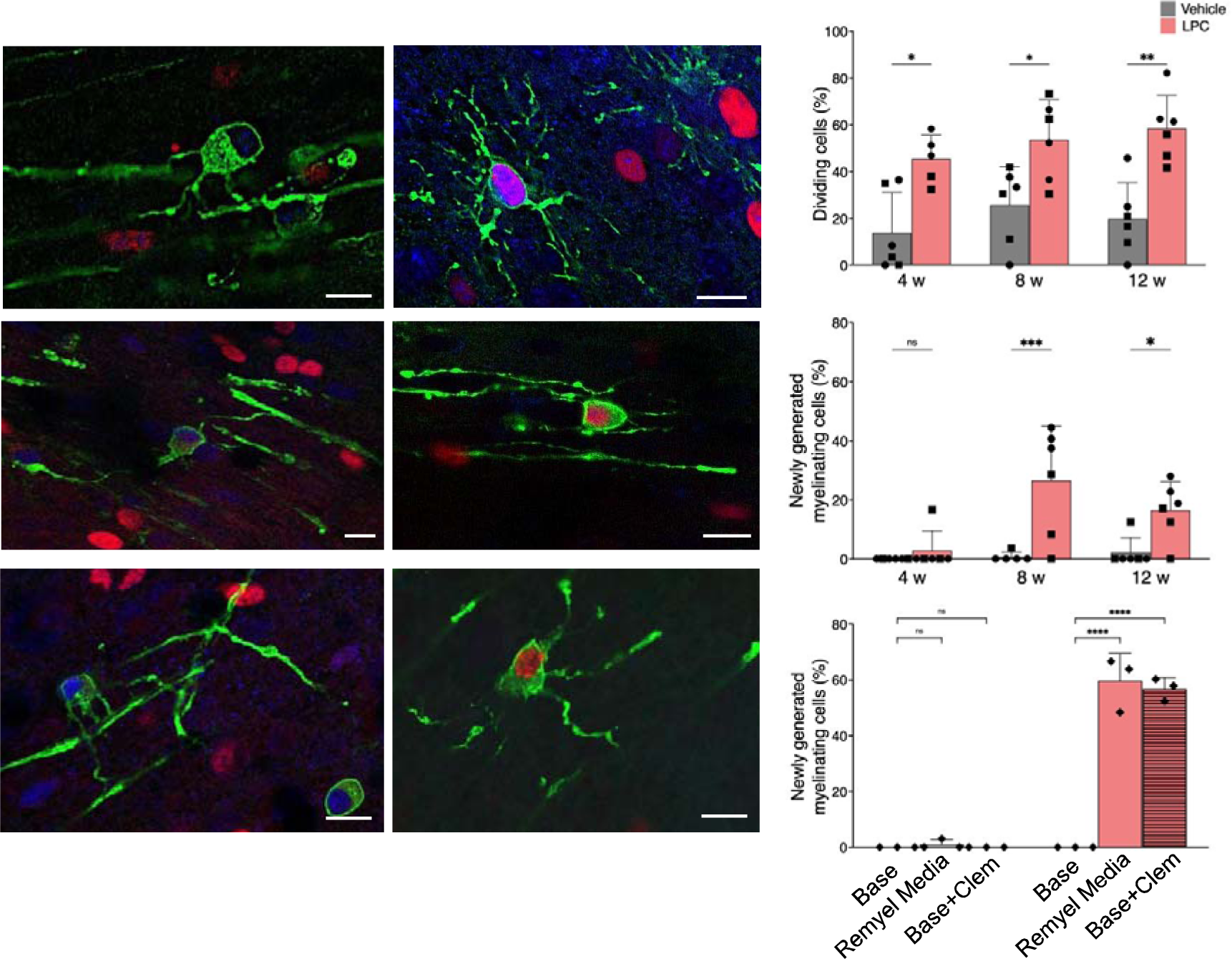
(a) CNP+SOX10+ oligodendrocytes co-expressing EdU are newly generated after LPC removal; scale bar = 20 μm. (b) The density of CNP+SOX10+EdU+ oligodendrocytes was higher in LPC treated organoids compared to Vehicle treated at all time points. (c) The density of CNP+SOX10+EdU+ myelinating oligodendrocytes as a proportion of all CNP+SOX10+EdU+ oligodendrocytes was significantly higher 8 w and 12 w after LPC removal. (d) Similarly to the remyelination media, clemastine treatment of organoids increased the density of myelinating CNP+SOX10+EdU+ oligodendrocytes compared to base media 8 w after LPC removal. Each data point depicts the sum from an individual organoid; different symbol depicts independent differentiations. Data graphed as mean ± SD and compared using a 2-way ANOVA. ns = not significant, * p<0.05 ** p<0.01 *** p<0.0005 **** p<0.0001

In summary, we have successfully generated iPSC-derived myelinating organoids that contain functional microglia and used these organoids to model key aspects of MS pathology. In response to an MS-relevant insult, myelin became fragmented and the resulting myelin debris was phagocytosed by microglia. We demonstrate that oligodendrocytes proliferate in response to myelin damage, differentiate and successfully generate new myelin, and this paradigm can be replicated with drug treatment. We therefore conclude that the generation of this model system has enormous potential to identify therapies that can promote new oligodendrocyte generation and new myelin formation.

## Acknowledgements

The authors acknowledge Prof Kevin Eggan, Prof Edouard Stanley, Dr Chris Bye, Dr Elizabeth Qian and Ms Katherine Lim for providing iPSC lines and the Florey Neuroscience Microscopy Facility and the Monash Biomedical Discovery Institute’s Imaging Platform for their support. We acknowledge Dr Magdaline Sakkas from the University of Melbourne Cytometry Platform for assistance with flow cytometry. We acknowledge Dr Ian Birchall for processing organoids for multiplex immunolabelling. This work was supported by a CASS Foundation Grant (S.B., D.G. & M.R.) and a National Health & Medical Research Council Ideas Grant (#2021811; D.G. & S.B.). S.B. was supported by a Rebecca L. Cooper Al & Val Rosenstrauss Medical Research Fellowship. B.T. was supported by FightMND Drug Screening Program and Stafford Fox Medical Research Foundation Grants. The schematics were created using Biorender.com.

## Online Methods

### Human iPSCs

The iPSC line 18a (female control line) was used for all assays and was kindly provided by Prof Kevin Eggan. The iPSC line C005 (female control line) was derived from donated fibroblasts in-house and reprogrammed using non-integrating vectors in feeder-free conditions following approval by the University of Melbourne Human Research Ethics Committee (#1749960.1). The iPSC line PB001-tdTomato (male control line) was kindly provided by Prof Ed Stanley. iPSCs were maintained on Matrigel (Corning)-coated 6 well plates in mTeSR1 (Stem Cell Technologies) at 37 °C and 5 % CO_2_ and passaged using EDTA (0.5 mM).

### Generation of iPSC-derived macrophage-like cells for seeding into organoids

Macrophage-like cells were generated from the PB001-tdTomato and 18a iPSC lines using adaptations to a previously published protocol [7]. The PB001-tdTomato derived cells were used for all data generation, unless otherwise stated. Briefly, there are two base medias used throughout; BASE I: 63:25:4 IMDM:F12:PFHMII (all Gibco), 0.25% Albucult (Albumedix), 0.15% polyvinyl alcohol (PVA; Sigma), 0.10% methylcellulose (Sigma), 1x ITS-X (Thermo), 50 µg/mL L-ascorbic acid 2-phosphate (AA2P; Sigma), 2 mM GlutaMAX (Thermo), 90 ng/mL Linolenic Acid (Sigma), 90 ng/mL Linoleic Acid (Sigma), 2.2 µg/mL SyntheChol (Sigma), 0.11 mM β-mercaptoethanol (B-ME; Sigma); BASE II is the same as BASE I but with PVA, methylcellulose and B-ME removed. At day 0, iPSCs were lifted into suspension and cultured in 10cm dishes in set-up media consisting of BASE I supplemented with 5 ng/mL Activin A (R&D Systems), 20 ng/mL BMP-4 (R&D Systems), 40 ng/mL VEGF (R&D Systems), 40 ng/mL SCF (R&D Systems), 5 ng/mL FGF-2 (Peprotech), 0.2 µM CHIR (StemCell Systems) and 10 µM Y-27632 (StemCell Technologies). On day 1, set-up media was replaced but with Activin A and Y-27632 removed and on day 2, set-up media was replaced but now also minus CHIR. On day 4, EBs were fed with maintenance media which is BASE I supplemented with 20 ng/mL BMP-4, 40 ng/mL VEGF, 40 ng/mL SCF, 10 ng/mL FGF-2, 25 ng/mL IL-3 (R&D Systems) and 50 ng/mL M-CSF (R&D Systems). From day 8, EBs were fed with maintenance II media (now with BASE II) minus BMP-4 and from day 11, EBs were fed with maintenance II media now supplemented with 50 ng/mL Flt3-Ligand (R&D Systems). At day 15, macrophage-like cells were collected; all single cells were extracted, strained (40 µm cell strainer; Corning), counted and frozen in liquid nitrogen until required for seeding into organoids.

### Generation of iPSC-derived Organoids

iPSC-derived organoids were generated from 18a and C005 as done previously [3] through adaptations to the original protocol [4]. 18a organoids were used for all data generation, unless otherwise stated. Briefly, iPSCs were split using EDTA (0.5 mM) at day -1 and on day 0 patterning commenced with media changed from mTeSR1 to phase I medium: 50:50 DMEM/F12+GlutaMAX:Neurobasal (both Gibco), 0.5 x B27 (Thermo), 0.5 x N2 (Thermo), 1 mM GlutaMAX (Thermo), 0.1 mM ascorbic acid (StemCell Technologies), 1 x anti-anti (Thermo), 1 mM N-acetyl cysteine (NAC; Sigma), 10 µM activin inhibitor (R&D Systems) and 100 nM LDN (Selleckchem). On day 7, cells were lifted using dispase into a T75 flask and maintained in phase II medium: 50:50 DMEM/F12+GlutaMAX:Neurobasal, 0.5 x B27, 0.5 x N2, 1 mM GlutaMAX, 0.1 mM ascorbic acid, 1 x anti-anti, 1 mM N-acetyl cysteine, 5 ng/mL FGF-2 (Peprotech), 0.1 µM retinoic acid (Sigma) and 5 ng/mL heparin (Sigma). On day 10-14 following neural rosette formation, cells were lifted using dispase into a low attachment 10 cm dish and maintained in phase III medium: Ad DMEM/F12 (Gibco), 1 x anti-anti, 0.5 x B27, 0.5 x N2, 1 mM glutaMAX, 5 ng/mL FGF-2, 1 µM purmorphamine (Sigma), 1 µM retinoic acid and 5 ng/mL heparin. After 7 days, media was changed to phase III-FGF: Ad DMEM/F12, 1 x anti-anti, 0.5 x B27, 0.5 x N2, 1 mM glutaMAX, 1 µM purmorphamine and 1 µM retinoic acid. After 14 days, media was changed to PRO media: Ad DMEM/F12, 1x anti-anti, 2 mM GlutaMAX, 1 x N2, 0.5 x B27, 5 µg/mL heparin, 10 ng/mL FGF-2, 20 ng/mL PDGF-AA (Peprotech), 0.5 µM purmorphamine, 10 ng/mL IGF-1 (Peprotech), 90 ng/mL T3 (Sigma) and 1 µM SAG (Merck). After 14 days, individual spheres were plated on PTFE-coated Millicell Cell Culture Inserts (Merck) to commence maturation and were therefore maintained in DIFF media: Ad DMEM/F12, 1x anti-anti, 2 mM GlutaMAX, 1 x N2, 0.5 x B27, 5 µg/mL heparin, 10 ng/mL IGF-1, 90 ng/mL T3 and 1x ITS-X (Thermo) thereafter; 16 organoids were plated per insert.

Three weeks after plate down onto inserts, 2000 macrophage-like cells were seeded per organoid and DIFF media was supplemented with IL-34 (100 ng/mL; R&D Systems) for 7 days to support integration, proliferation and differentiation.

### Validation of differentiation of macrophage-like cells to microglia within our organoids

After 11 weeks in DIFF media (and therefore eight weeks following macrophage seeding), organoids were lifted and digested into single cells using papain (Worthington; following kit protocol). 300,000 cells (per antibody) were suspended in FACS buffer (100 µL 0.5% BSA + 1mM EDTA in PBS minus Ca^2+^ and Mg^2+^) and then incubated with CD45 (FITC Clone 2D1 #368507; 5 µg/mL), CD11b (APC clone ICRF44 #982604; 10 µg/mL), P2RY12 (APC clone S16001E #392113; 20 µg/mL) and CX3CR1 (647 clone 2A9-1 #341607; 5 µg/mL) for 30 min on ice. Cells were spun at 300 x g for 3 min and washed twice with FACS buffer. Propidium iodide (1 µg/mL) was added to the cells to aid in elimination of dead cells prior to data acquisition. Cells were run to acquire FSC-H vs SSC-H followed by FSC-H vs FSC-A for doublet discrimination. Cells containing no other markers (other than the intrinsic tdTomato positive cells) was run to establish the gate for all tdTomato positive cells. Dissociated organoids with microglia with no intrinsic reporter (organoids with 18a microglia) were also run to confirm the accuracy of the gate. Frozen 3 week tdTomato+ macrophage-like cells were thawed following the same protocol as when the cells were seeded into organoids and was stained with CD45 and run to establish shift in phenotype between pre-seeded cells and cells cultured for 11 weeks in organoids.

### Lysolecithin Treatment

After 11 weeks in DIFF media, lysolecithin (0.1 mg/mL; Sigma; LPC) was added (or ethanol v/v as vehicle control; Vehicle). All media was removed from underneath the inserts and 1 mL of DIFF+LPC/Vehicle was added below the insert and 1 mL of DIFF+LPC/Vehicle was added to the top of the insert. Plates were placed back in the incubator and left for 16 h. All media was then aspirated from above and below the insert and fresh DIFF media was added below the insert; organoids were then cultured as normal. 5 h, 24 h, 7 d, or 14 d after LPC/Vehicle removal, organoids were fixed in 4% PFA for 2 h. LPC has been extensively used *in vivo* and in rodent slice cultures is typically used at 0.5 mg/mL [15, 16]; it has been used in iPSC-derived organoids at a concentration of 0.25 mg/mL [17].

### Live imaging of tdTomato-positive microglia within organoids

To visualise movement and activity of tdTomato+ microglia within organoids, 11 week old organoids were transferred to 6-well glass bottom plates with #1.5 cover glass (Cellvis) for live-imaging on an inverted CSU-W1 T2 Marianis spinning disk confocal (3i) using a 50 μm disk, and a LD C-Apo 40x/1.1 Water CORR objective lens. Tdtomato was excited with 561nm laser light, and emission light was captured on a BSI Express air-cooled sCMOS camera. Temperature, CO_2_ and humidity were maintained at 37 °C, 5 %, and 80 % respectively. 20 μm z-stacks were obtained at each time point, which were maximum intensity projected in post processing to produce the videos presented. Baseline activity was captured at 12 s intervals for 120 min. To stimulate a microglial response to light damage, organoids were photostimulated using the Phasor 1-photon 8T holographic photostimulation system, in a region directed next to cells within the field of view.

To witness whether microglia responded directly to LPC, 3 organoids from one differentiation from each condition were imaged immediately after the introduction of Vehicle or 0.1 mg/mL LPC, with each organoid imaged at an interval of 4 min for a duration of 16 h. Seven days after LPC (or vehicle) removal, 3 separate organoids per condition from one differentiation were imaged at an interval of 5.35 s for a duration of 2 min 14 s.

### Remyelination Assay

Remyelination Media: 24 h after LPC/Vehicle removal, some inserts commenced the remyelination assay. Organoids were fed with DIFF media supplemented with BDNF (20 ng/mL; Peprotech), NT-3 (20 ng/mL; R&D Systems), PDGF-AA (10 ng/mL) and IL-34 (100 ng/mL) until 7 d after which organoids were fed with just DIFF media. Throughout the remyelination assay, media was also supplemented with EdU (1 µM; Thermo). EdU is incorporated into newly synthesised DNA in dividing cells and is therefore absent from post-mitotic oligodendrocytes that survived the demyelinating injury. After 4 w, 8 w and 12 w, organoids were fixed in 4 % PFA for 2 h.

Clemastine: 24 h after LPC/Vehicle removal, membranes were randomised to either Base media, Remyelination media or Base+Clemastine. Base organoids were maintained in Ad DMEM/F12, 1x anti-anti, 2 mM GlutaMAX, 1 x N2, 0.5 x B27 and 5 µg/mL heparin. Remyelination organoids were treated as above. Clemastine organoids were maintained in Base media supplemented with 1 µM Clemastine Fumarate for 8 w (Sigma). All medias were supplemented with 1 µM EdU and organoids were fixed at 8 w in 4 % PFA for 2 h.

### Immunostaining for assessment of demyelination, microglial morphology, myelin phagocytosis, new oligodendrocyte generation and new myelin generation

Organoids were immunolabelled whole; individual organoids (4-6 per analyses) were transferred into a 96 well plate and washed three times in 0.2 M PBS. Organoids were permeabilised (40 min in 0.3 % PBS-TritonX) and blocked (2 h in 10 % NDS in 0.3% PBS-TritonX) after which they were incubated overnight at 4 °C in primary antibodies in blocking buffer (myelin basic protein; MBP, 1:1000, MAB7349 and Iba-1, 1:300, 019-19741). For the new oligodendrocyte generation, after the initial block organoids underwent antigen retrieval for 20 min at 99 °C in citrate buffer (pH 6.0) followed by another blocking step for 2 h after which they were incubated in primary antibodies (2’,3’-cyclic nucleotide 3’ phosphodiesterase; CNP, 1:2000, AMAB91072 and SOX10, 1:300, Ab155279) in blocking buffer overnight at 4 °C.

Organoids were washed three times in 0.1 % PBS-Tween20 followed by incubation with secondary antibodies in blocking buffer (MBP in 488, Iba-1 in 647, CNP in 488 and SOX10 in 405) for 2 h at room temperature. Organoids were washed in PBS. The MBP/Iba-1 organoids were incubated in Hoescht (1:1000; Thermo) for 10 min prior to washing in PBS and mounted onto slides using Dako Mounting Medium. The CNP/SOX10 organoids were labelled for EdU (C10340) according to kit instructions post secondary antibodies after which they were washed and mounted onto slides using Dako Mounting Medium. All slides were blinded prior to imaging.

### Imaging and Analysis for assessment of demyelination, microglial morphology, myelin phagocytosis, new oligodendrocyte generation and new myelin generation

Organoids were imaged using a Zeiss 780 Laser Scanning Confocal microscope (Carl Zeiss AG, Oberkochen, Germany) and acquired with Zen Black (Zen 2012 SP5, version 14.0.23.201). For the demyelination, microglial morphology and myelin phagocytosis analyses, three organoids were imaged per time point per differentiation (3-4 independent differentiations for 18a; 1 independent differentiation for C005) and 4 z-stacks (30 µm z-stack, z-step size of 1 µm) were taken per organoid at consistent positions using a Plan Apo 20x/0.8 NA DIC objective. For the new oligodendrocyte generation, 4-8 organoids were imaged per time point per differentiation (5 independent differentiations at 4 w and 4 independent diffs at 8 and 12 w). For newly myelinating oligodendrocytes, 4-8 organoids were imaged per time point per differentiation (4 independent differentiations of which 2 had measurable myelin remaining in the organoids, irrespective of treatment group, which is as expected given the long culturing time). For both analyses, 4-12 images were taken using a Plan Apo 40x/1.4 DIC or a Plan Apo 63x/1.4 Oil DIC objective (z-stack size also dependent on spanning of myelin sheaths) and were taken from positions of positive CNP staining.

For the demyelination, microglial morphology and myelin phagocytosis analyses, Imaris software (Bitplane, version 9.9.1) was used. For the demyelination analyses, the MBP channel for each image was loaded; a surface mask was applied to label each myelin segment and a bounding box threshold of 10 µm was applied as an attempt to limit the inclusion of MBP+ pre-myelinating oligodendrocytes. The average sheath length was then calculated per field of view, averaged per organoid and each individual organoid constitutes one data point. For the C005 iPSC-derived organoids where the myelination is less extensive, the average sheath length was calculated per field of view and each field of view constitutes one data point. For the microglial morphology analyses, the Iba-1 channel for each image was loaded and a surface mask was applied to label all individual microglia. An additional filament mask was applied to the surface mask to calculate the cellular parameters such as cell soma area and filament area. The average cell soma area (representing the cell body) and cell filament area (representing the microglial processes) were calculated per field of view, averaged per organoid and each individual organoid constitutes one data point. For the myelin phagocytosis analyses, both MBP and Iba-1 channels for each image were loaded and a surface mask was applied to both the channels. The distance between the surfaces was set to - 1 μm to find the inclusion of MBP+ myelin within Iba-1+ microglia and the percentage of phagocytosis was calculated based on the percentage of Iba-1+ area occupied by MBP+ debris. For the representative images in Fig 2e and Suppl Fig 5a, the surface-surface coloc function Xtension was applied to visualise phagocytosis (Imaris version 9.9.1).

For the new oligodendrocyte generation analyses, the cell counter plug-in in Fiji was used. All CNP+SOX10+ cells were quantified per field of view (through entire z-stack) and summed per organoid then the proportion of those cells that were triple positive for EdU+ were counted and summed per organoid to ascertain the proportion of all oligodendrocytes that were newly generated since the LPC/Vehicle insult. To determine which of those newly generated oligodendrocytes were myelinating, additional analyses were completed. Of the CNP+SOX10+EdU+ oligodendrocytes, the proportion that had commenced myelinating was calculated and were classified as such if CNP+ tubular structures were present attached to the cell body via a cytoplasmic process. For both measurements, all images were summed to calculate the proportion per organoid and each data point constitutes one organoid.

### Microglial Phenotyping Using Multiplex Immunolabelling

Multiplex immunolabelling and analyses were carried out as done previously [18]. Organoids (6-8 per time point) for 1-2 independent differentiations were processed following fixation (20 min 70% EtOH, 20 min 95% EtOH, 20 min 95% EtOH, 20 min 100% EtOH, 20 min 100% EtOH, 20 min Xylene, 1 h Paraplast+ wax) followed by embedding in Paraplast+ wax. Blocks were sectioned (7 µm) onto ÜberFrost® Printer Slides (InstrumeC); two sections per slide. Slides were cleared and rehydrated prior to antigen retrieval in Tris-EDTA with 0.05% Tween20 (pH 9.0) followed by autofluorescence quenching with TrueBlack and incubation with primary antibodies in a 1% NGS buffer overnight at 4 °C (L-ferritin 1:1000 Ab218400, HLA-DR 1:1000 M0775, CD68 1:200 Ab783, Iba-1 1:1000 #234-004). Slides were washed and incubated with relevant AlexaFluor secondary antibodies (1:500) for 4 h at room temperature after which they were incubated in Hoechst 33258 and coverslipped using ProLong Gold Antifade mounting media (Thermo).

Slides were imaged on a Zeiss Z2 Axioimager (20x/0.9 NA) using MetaSystems VSlide acquisition software and MetaCyte stitching software. 2-3 organoids were analysed per marker, per treatment, per differentiation. We developed custom image analysis pipelines in Metamorph software (Molecular Devices) similar to those previously described [18, 19]. Two image analysis pipelines were developed for the assessment of microglial functional marker levels: tissue-wide and single-cell analyses. Tissue-wide analyses were used to assess whether total microglial expression of microglial markers was altered with LPC treatment. The tissue-wide analysis quantified the average labelling intensity of each microglial marker across all Iba-1-positive microglia in the iPSC-derived organoids. To identify all Iba1-positive cells, the Iba-1 image was first edited using the Background Correct tool before all cells were identified using the Adaptive Threshold tool generating a binary mask. Within this binary mask, the average labelling intensity of L-ferritin, HLA-DR, CD68, and Iba-1 was measured. These average intensity measures represent the average protein expression level across the microglia in each iPSC-derived organoid. The single-cell analysis quantified the average labelling of each microglial marker within each Iba-1-positive microglia in the iPSC-derived organoids. The Iba-1-positive binary mask was generated as per the tissue-wide analysis above. Each object within this binary mask was considered an Iba-1-positive microglia, and the average intensities of L-ferritin, HLA-DR, CD68, and Iba-1 were measured within each microglia. These average intensity measures represent the average protein expression in each microglia.

### Statistical Analysis

All data are expressed as mean ± standard deviation with individual symbols depicting average data from one organoid, unless otherwise stated. Intergroup differences were appropriately assessed by using a 2-way ANOVA with Šidák’s multiple comparisons post-hoc test with the two variables being treatment and time. The microglial single-cell average intensities were compared between LPC and Vehicle treated organoids using unpaired t-tests. Statistical analysis were performed on Prism 9 (GraphPad).

## Supplemental Figures

**Supplemental Figure 1:**
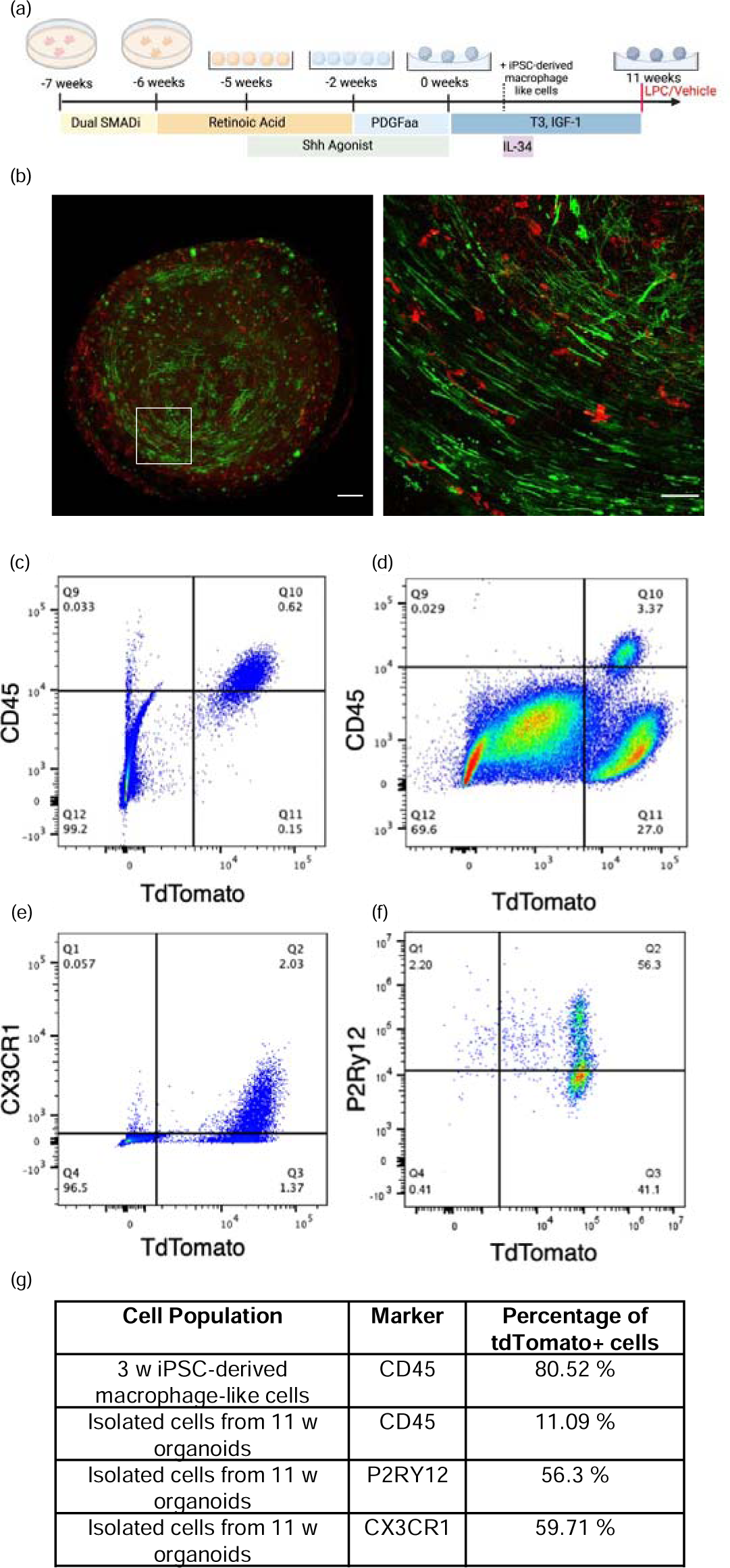
(a) Schematic of protocol for generating myelinating organoids; our protocol consists of 7 weeks of patterning followed by 11 weeks of maturation (the latter is when myelination occurs). Organoids were seeded with iPSC-derived macrophage-like cells at 3 weeks maturation (dotted line) and LPC/Vehicle was added at 11 weeks maturation (red line). (b) A representative tiled image (a 5×5 tile taken using a Plan Apo 10x/0.45 DIC objective at 1.2 x zoom) of one z-plane through an entire organoid at 11 weeks maturation co-labelled for MBP (green) and Iba-1 (red); scale bar = 200 μm. White box depicts the inset image (a maximum projection image of a 22 μm z-stack with 1 μm steps taken using a Plan Apo 20x/0.8 DIC objective); scale bar = 50 μm. Single cell suspensions from 3 week iPSC-derived tdTomato+ macrophage-like cells were stained with CD45. Single cell suspensions from 11 week organoids containing tdtTomato+ cells were stained with immune markers: (c) CD45; (d) CX3CR1; and (e) P2RY12. (f) quantification of the proportion of tdTomato+ cells

**Supplemental Figure 2:**
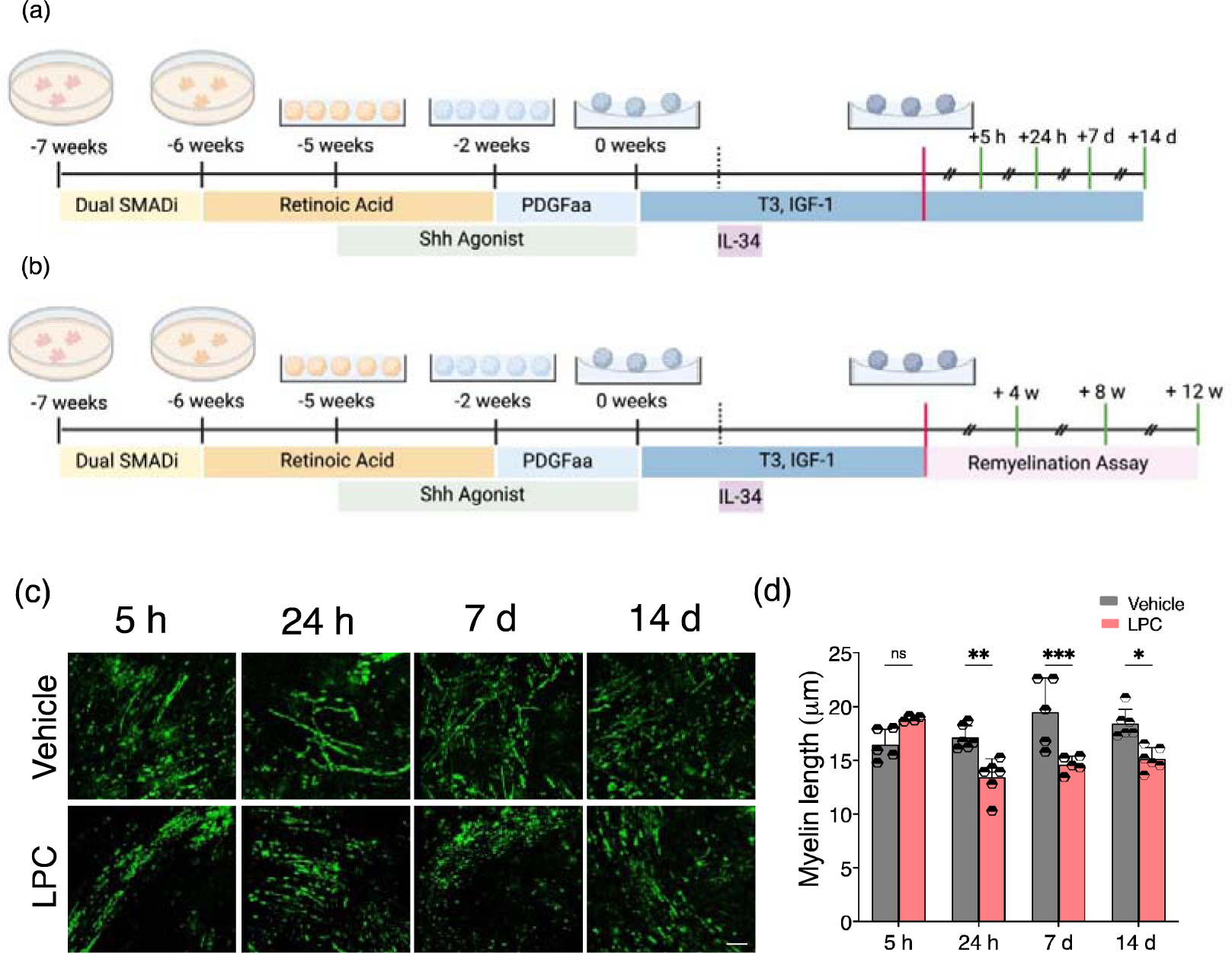
Schematic of protocols with time points at which organoids were collected after LPC/Vehicle (red line) for assessment of (a) demyelination and microglial morphology and function (5 h, 24 h, 7 d and 14 d) and (b) new oligodendrocyte and myelin formation (4 w, 8 w and 12 w) (green lines). (c) In organoids derived from the C005 iPSC line, LPC triggered overt MBP-positive myelin fragmentation compared to Vehicle treated organoids; scale bar = 50 μm. (d) Average myelin length per field of view in C005 derived organoids was significantly shorter in LPC treated organoids compared to Vehicle treated, at 24 h, 7 d and 14 d. Each data point depicts the average from an individual field of view; different symbols depict independent differentiations. Data graphed as mean ± SD and compared using a 2-way ANOVA. ns = not significant, * p<0.05 ** p<0.01 *** p<0.0005 **** p<0.0001

**Supplemental Figure 3:**
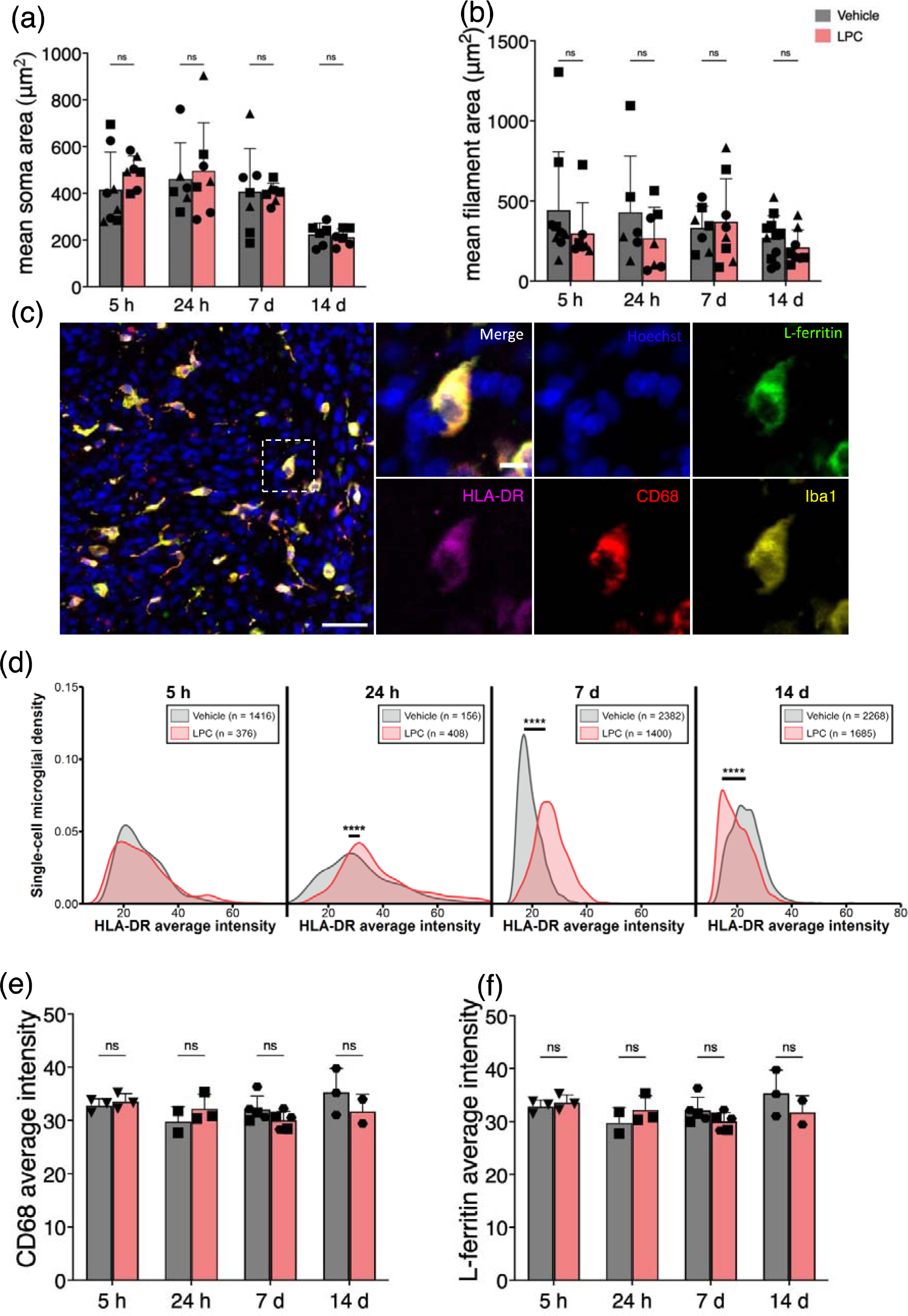
(a) mean soma area and (b) mean filament area, as a surrogate measure for microglial cell body and process area, respectively, were not different between groups at any time point. (c) multiplex immunolabelling was used to further stratify the microglial population; scale bar = 50 μm (10 μm in inset image highlighted by white box). (c) single cell analysis of the average intensity of HLA-DR expression in Iba-1+ microglia showed unimodal expression at all time points with significant differences in expression at 24 h, 7 d and 14 d. Tissue wide average intensity of (e) CD68 and (f) L-ferritin in Iba-1+ microglia was not different between groups at any time point. Each data point depicts the average from an individual organoid; different symbols depict independent differentiations. Data graphed as mean ± SD and compared using a 2-way ANOVA (single cell data in (d) compared using unpaired t-tests). ns = not significant, **** p<0.0001.

**Supplemental Figure 4:**
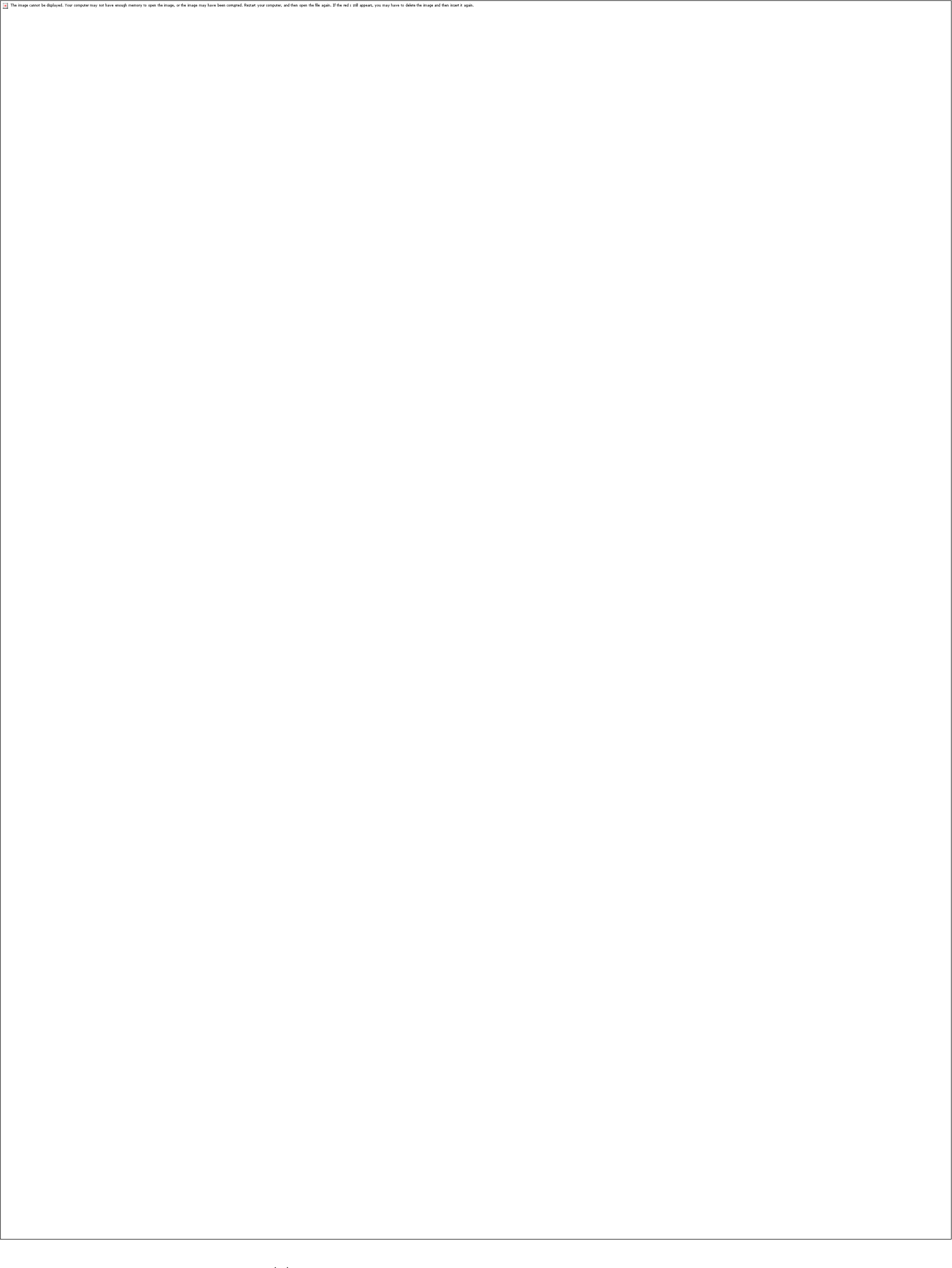
(a) Violin plots depicting single cell average intensity of Iba-1, HLA-DR, CD68 and L-ferritin expression within Iba-1+ microglia at 5 h, 24 h, 7 d and 14 d after LPC removal. At 5 h, LPC (3 organoids) and Vehicle (3 organoids) were analysed from one independent differentiation. At 24 h, LPC (3 organoids) and Vehicle (2 organoids) were analysed from one independent differentiation. At 7 d, LPC (5 organoids) and Vehicle (5 organoids) were analysed from two independent differentiations. At 14 d, LPC (2 organoids) and Vehicle (2 organoids) were analysed from one independent differentiation. (b) heatmap to depict percentage change in marker expression in LPC treated organoids relative to Vehicle.

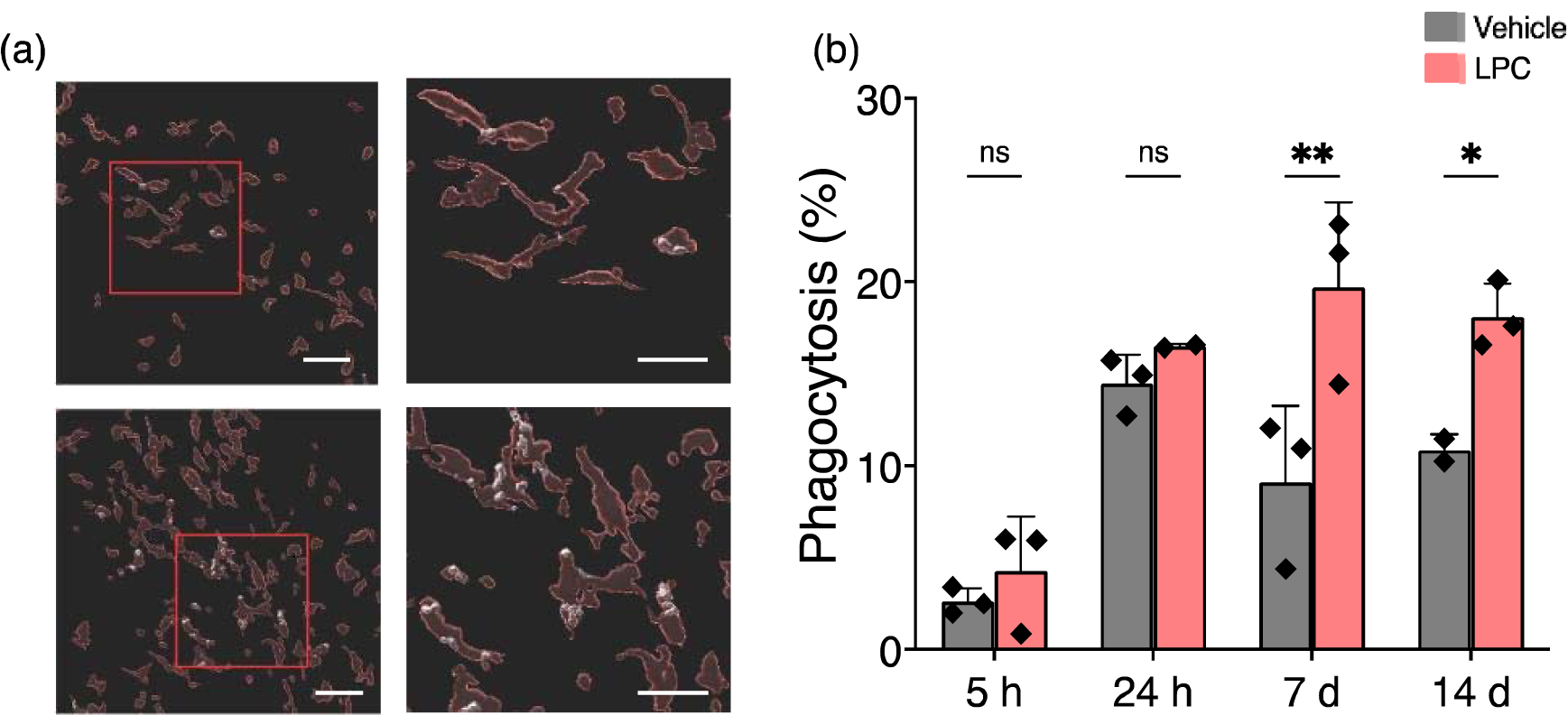

**Supplemental Video 1:** live imaging of tdTomato+ microglia in an untreated 11 week organoid. Over 80 min, microglia extend and retract ramified processes, ostensibly surveying the local environment, with some cells migrating small distances; scale bar = 50 μm.

**Supplemental Video 2:** live imaging of tdTomato+ microglia in an untreated 11 week organoid, before and after focal photostimulation with Phaser laser; boxes within the field of view represent positions of Phaser laser. In the initial 5 min after laser, cells retract cellular processes, but within 15 min they have extended their processes towards the phaser laser locations. Scale bar = 50 μm.

**Supplemental Video 3:** live imaging of tdTomato+ microglia for 16 h in an 11 week organoid immediately (t=0) following vehicle supplementation into media; scale bar = 50 μm.

**Supplemental Video 4:** live imaging of tdTomato+ microglia for 16 h in an 11 week organoid immediately (t=0) following LPC supplementation into media; scale bar = 50 μm.

**Supplemental Video 5:** live imaging of tdTomato+ microglia in an organoid 7 d after vehicle removal; scale bar = 50 μm.

**Supplemental Video 6:** live imaging of tdTomato+ microglia in an organoid 7 d after LPC removal; scale bar = 50 μm.

